# Early overactivation of non-muscle myosin II during adaptation to combined BRAF and MEK inhibitors in dedifferentiating cutaneous melanomas

**DOI:** 10.64898/2025.12.03.686756

**Authors:** Andrea Garcia-Perez, Lucía Sanchez-Garcia, Marta Duran-Renieblas, Federica Sella, Silvia Novo-Acedo, Angel Delgado-Lopez, Adrian Barreno, Judith Gracia, Christopher Rios, Erica J. Young, Laszlo Radnai, Clara Diaz-Utrilla, Julia M. Martinez-Gomez, Rosa M. Marti, Marta C. Sallan, Oscar Maiques, Courtney A. Miller, Eva Perez-Guijarro, Anna Macià, Mitchell P. Levesque, Jose L. Orgaz

**Affiliations:** Universidad Carlos III de Madrid, Departamento de Neurociencia y Ciencias Biomédicas, 28903 Getafe, Madrid, Spain; Instituto de Investigaciones Biomédicas Sols-Morreale (IIBM), Consejo Superior de Investigaciones Científicas-Universidad Autónoma de Madrid, 28029 Madrid, Spain; Department of Dermatology, University of Zurich, University Hospital Zurich, 8952 Schlieren, Switzerland; Oncologic Pathology Group, Institut de Recerca Biomèdica de Lleida (IRBLleida), University of Lleida, 25198 Lleida, Spain; Department of Molecular Medicine and Department of Neuroscience, The Herbert Wertheim UF Scripps Institute for Biomedical Innovation & Technology, Jupiter, FL 33458, USA; Department of Biochemistry, Faculty of Medicine, Universidad Autónoma de Madrid (UAM), 28029 Madrid, Spain; Barts Cancer Institute, Queen Mary University of London, Charterhouse Square, London, United Kingdom; Cancer Biomarkers & Biotherapeutics, Barts Cancer Institute, Queen Mary University of London, London, United Kingdom; Centre of Biomedical Research on Cancer (CIBERONC), Instituto de Salud Carlos III (ISCIII), 28029 Madrid, Spain; Department of Dermatology, Hospital Universitari Arnau de Vilanova de Lleida, Institut de Recerca Biomèdica de Lleida (IRBLleida), University of Lleida, 25198 Lleida, Spain; Instituto de Investigación Sanitaria Hospital Universitario La Paz (IdiPAZ), 28046 Madrid, Spain

**Keywords:** melanoma, therapy tolerance, resistance, myosin, cytoskeleton

## Abstract

Cutaneous melanoma is a very aggressive type of skin cancer with remarkable phenotypic plasticity that contributes to adaptation and resistance to targeted therapies against the MAPK pathway. Previous research described that non-muscle myosin II (NMII) of the actomyosin cytoskeleton, which is essential for cell migration and metastasis, is overactivated in BRAF inhibitor-resistant melanomas. Since the combination of BRAF and MEK inhibitors (BMi) is the current standard of care, we investigated if and how NMII activity is regulated during adaptation to BMi. Here, we find that most dedifferentiating BMi-resistant melanomas overactivate NMII compared to their parental counterparts. NMII activity generally increases during the first 2 weeks of BMi treatment, and it is followed by elevated total NMII levels due partly to transcriptional modulation. Although ERK activity rebounds with similar kinetics, NMII overactivation is not prevented by ERK inhibition but by blockade of ROCK. In melanomas that hyperdifferentiate during adaptation to BMi, NMII activity is not increased upon BMi treatment due, in part, to MITF. We also find that co-targeting NMII along BMi in some melanomas reduces survival of drug-tolerant persister cells, which would delay the development of resistance. Therefore, our study identifies elevated NMII activity as a potential marker of adaptation to MAPK in some melanoma subpopulations, and also provide an approach to delay the emergence of resistance to MAPK-targeted therapy.

## Introduction

Inhibition of ERK signalling in the MAPK pathway using BRAF inhibitors (BRAFi) (1,2), and later with the combination of BRAFi and MEK inhibitors (MEKi) (3–5), reduces tumour growth and prolongs survival of most patients with metastatic BRAF^V600E^ cutaneous melanomas. However, most cases develop resistance to these therapies, which complicates patient outcomes (3–8).

Resistance to MAPK inhibitors (MAPKi) involves genetic (e.g., mutations) and non-genetic changes that lead to extensive epigenetic, transcriptional and translational re-wiring (9–14). These events are key especially during early adaptation to therapy, contributing to survival of residual drug-tolerant persister cells (DTPs) (15,16). DTPs survive the initial therapy challenge and remain almost quiescent (17) in this reversible, transient state (15,18). Eventually, DTPs resume proliferation as drug-tolerant proliferating persister cells (DTPP) (14), evolving to a more stable, acquired resistant state (12,13,19) responsible for tumour re-growth and patient relapse (11).

Upon MAPKi treatment of a drug-naïve melanoma with high/moderate sensitivity to MAPKi (melanocytic and differentiated phenotype), some subpopulations undergo a phenotype switching that enables survival (10–13,20,21). Cells can proceed through diverging trajectories towards stable therapy-resistant states, by either switching to an hyperdifferentiated/pigmented state, or to more dedifferentiated states (neural crest stem cell (NCSC) or the most dedifferentiated mesenchymal/invasive) (10–13,20,22,23).

Some of these phenotypes are also found in therapy-naïve melanomas (melanocytic, intermediate/transitory, NCSC, dedifferentiated/mesenchymal/invasive); not surprisingly, the mesenchymal/invasive phenotype displays intrinsic resistance to BRAFi (21). In turn, BRAFi-resistant melanomas are generally more invasive and metastatic (24–29). Increased invasion and metastasis usually involves activation of the actin and non-muscle myosin II (NMII) or actomyosin cytoskeleton (30,31).

NMII is a holoenzyme comprised of several heavy and light chains. NMII activity is regulated by phosphorylation events, in particular in residues within its regulatory light chain (MLC2) by kinases Rho-kinase (ROCK) and myosin light chain kinase (MLCK), among others (31,32). Ser19 phosphorylation of MLC2 (p-MLC2) induces a conformational change that extends NMII (active conformation) and increases its actin-binding/ATPase activity (33). An additional phosphorylation of Thr18 (pp-MLC2) stabilises NMII in the active conformation and further controls contraction by modulating the ATPase activity (34) and the affinity of the complex for actin to assemble into actomyosin filaments (32), which generate contractile force that drives cell migration, among other processes (31).

Both DTP and BRAFi-resistant cells display extensive actomyosin cytoskeleton remodelling due to activation of upstream regulators, like Rho GTPases and their effectors, and focal adhesion signalling linking to the extracellular matrix (24,35). Furthermore, there is mounting evidence supporting a wider role of cytoskeletal remodelling under therapeutic pressure, since DTP/resistant cells become more dependent on the cytoskeleton and mechanical signalling for survival upon treatment (10,36–39). BRAFi-resistant melanomas have increased NMII activity levels compared to their parental counterparts (29,38,39), creating a dependency in resistant cells for NMII regulator ROCK. This vulnerability can be therapeutically exploited by co-targeting ROCK to overcome resistance to BRAFi (38,39).

Since the current standard of care is the combination of BRAFi and MEKi (BMi), in this study we sought to elucidate if and how NMII activity is regulated during adaptation to BMi. Most BMi-resistant (BMR) melanomas with a more dedifferentiated phenotype increase NMII activity, and this occurs early on during adaptation to BMi. In several melanoma models combination of NMII inhibitors (NMIIi) with BMi delays emergence of persister cells. Taken together, our study identifies high NMII activity as a biomarker of early adaptation to therapy in most melanomas, and the potential of a combination therapy with NMIIi to delay resistance.

## Results

### BRAFi+MEKi-Resistant melanoma sub-lines have higher levels of NMII activity than their parental counterparts

We generated 7 BMR sub-lines after chronic exposure to BMi (dabrafenib+trametinib) of 5 BRAF^V600E^ cutaneous melanoma cell lines and 2 patient-derived short-term melanoma cultures (MMxxx) (40,41) (Fig. 1a, S1a-b). All 7 BMR sub-lines had increased NMII activity (pp-MLC2) and, in some cases, elevated total MLC2 levels compared to parental counterparts (P). Immunofluorescence analyses confirmed the increased NMII activity (pp-MLC2 and p-MLC2) and actin fibres in BMR sub-lines (Fig. 1b, S1c).

**Figure 1.**
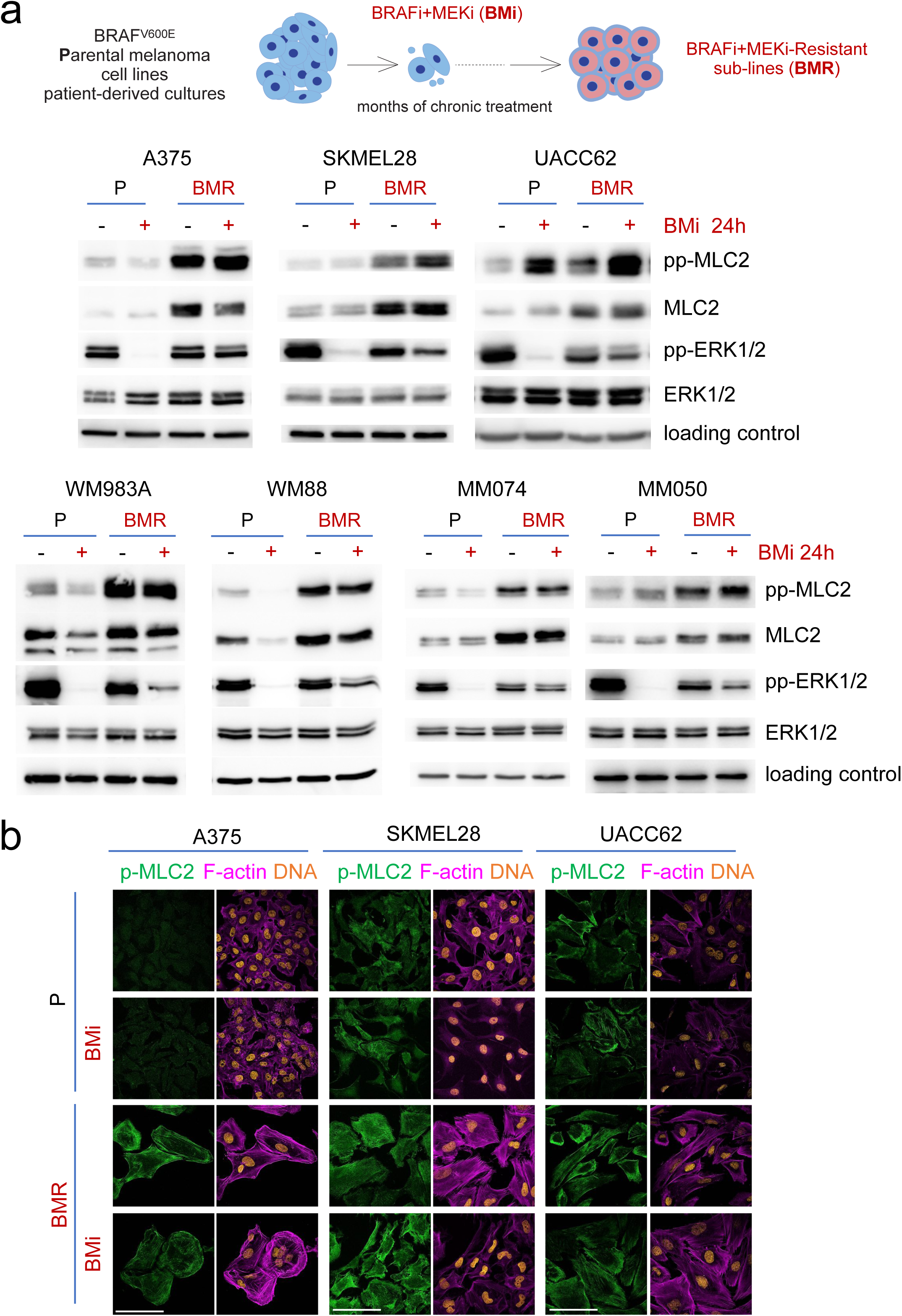
BRAFi+MEKi-resistant melanoma sub-lines have higher levels of NMII activity than their parental counterparts. (**a**) Top, schematic of generation of BRAFi+MEKi (BMi)-resistant (BMR) sub-lines. Bottom, immunoblots of indicated proteins from pairs of parental (P) and BMR lines after 24 h treatment with BMi (125 nM BRAFi dabrafenib + 6.25 nM MEKi trametinib; except WM983A (1 μM + 50 nM), WM88 (0.5 μM + 25 nM)). Loading control was α-tubulin except for WM983A and MM074 that had RhoGDI. (**b**) Confocal immunofluorescence images of p-MLC2, F-actin (phalloidin) and DNA (DAPI) stainings after 24 h treatment with BMi. Scale bars, 100 µm.

Many BRAFi-R and BMR melanomas are more dedifferentiated than their corresponding baseline/parental (P) (10,12,14,20,21). We tested levels of phenotype markers AXL, NGFR, SOX10 and MITF (12,21,27,41,42) to phenotypically characterise these models (Fig. S1b). All 7 BMR sub-lines were more dedifferentiated compared to P lines, even though P lines differed in their phenotype (e.g., A375 was NCSC (41), while SKMEL28 was melanocytic). Therefore, NMII activity increases in dedifferentiated BMR melanomas, regardless of the phenotype of the parental cell line.

### NMII levels increase early on during adaptation to BRAFi+MEKi

Next, we assessed how early NMII activity increases during adaptation to BMi. We treated P lines with BMi and harvested cell lysates at different time points. In A375 cells, pp-MLC2 increased after 2-3 days of BMi treatment compared to control (Fig. 2a-b), while total MLC2 levels augmented after 6-9 days of treatment. We obtained similar results with other approved BMi (encorafenib+binimetinib, Fig. S2a).

**Figure 2.**
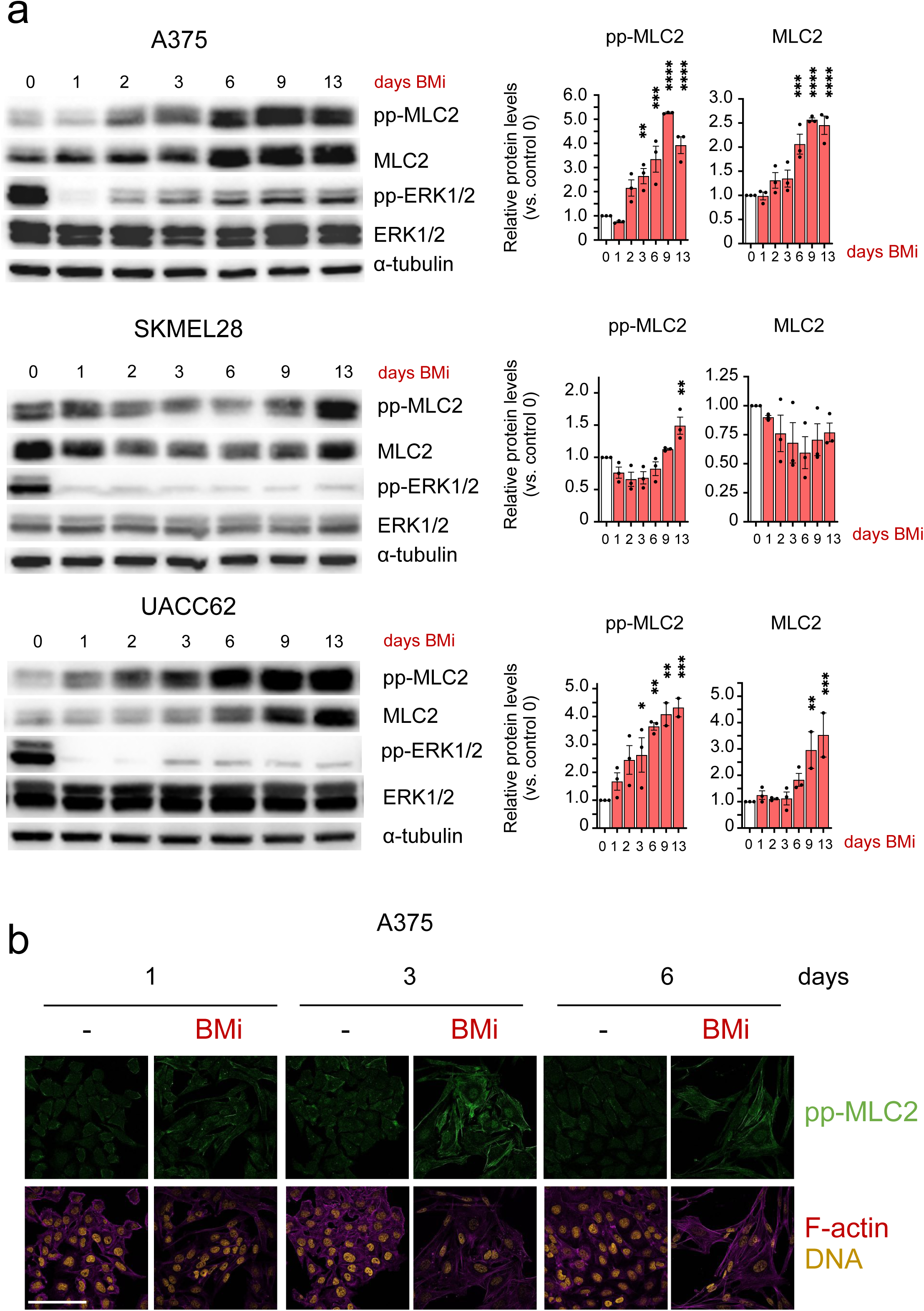
NMII levels increase early on during adaptation to BRAFi+MEKi. (**a**) Immunoblots of indicated proteins from parental melanoma cell lines after treatments with BMi (125 nM BRAFi dabrafenib + 6.25 nM MEKi trametinib). Representative of 3 independent experiments. Graphs show relative quantification vs DMSO control (0). P-values of comparisons of each day BMi vs DMSO control by one-way ANOVA with Dunnett post hoc test (*p<0.05, **p<0.01, ***p<0.001, ****p<0.0001). (**b**) Confocal immunofluorescence images of pp-MLC2, F-actin (phalloidin) and DNA (DAPI) stainings of A375 cells after treatment with BMi. Scale bar, 100 µm.

In other cell lines, NMII activity displayed slightly different kinetics, with pp-MLC2 levels increasing after 2-3 days (UACC62, WM983A), 13 days (SKMEL28) or just 1 day (mouse melanoma YUMM1.7) of BMi treatment (Fig. 2a, S2a). Similar to the human BMR, YUMM1.7.BMR cells displayed increased pp-MLC2 levels compared to P cells (Fig. S2a).

We also analysed levels of the phenotype markers during adaptation to BMi. While in A375 and SKMEL28 cells levels of markers were modulated at similar time points as pp-MLC2, in UACC62 cells the increase in pp-MLC2 levels preceded those of AXL and NGFR (Fig. S2b). These data suggest that, in some models, the dynamics of NMII activity may not be linked to the phenotype switching during adaptation to BMi.

In order to evaluate the preclinical relevance of our findings, we quantified NMII activity by immunohistochemistry in tumour samples from YUMM1.7 mouse allografts treated with BRAFi (Fig. 3). We found that p-MLC2 levels increased after 3 days of BRAFi in vivo, validating our previous results in cell line models.

**Figure 3.**
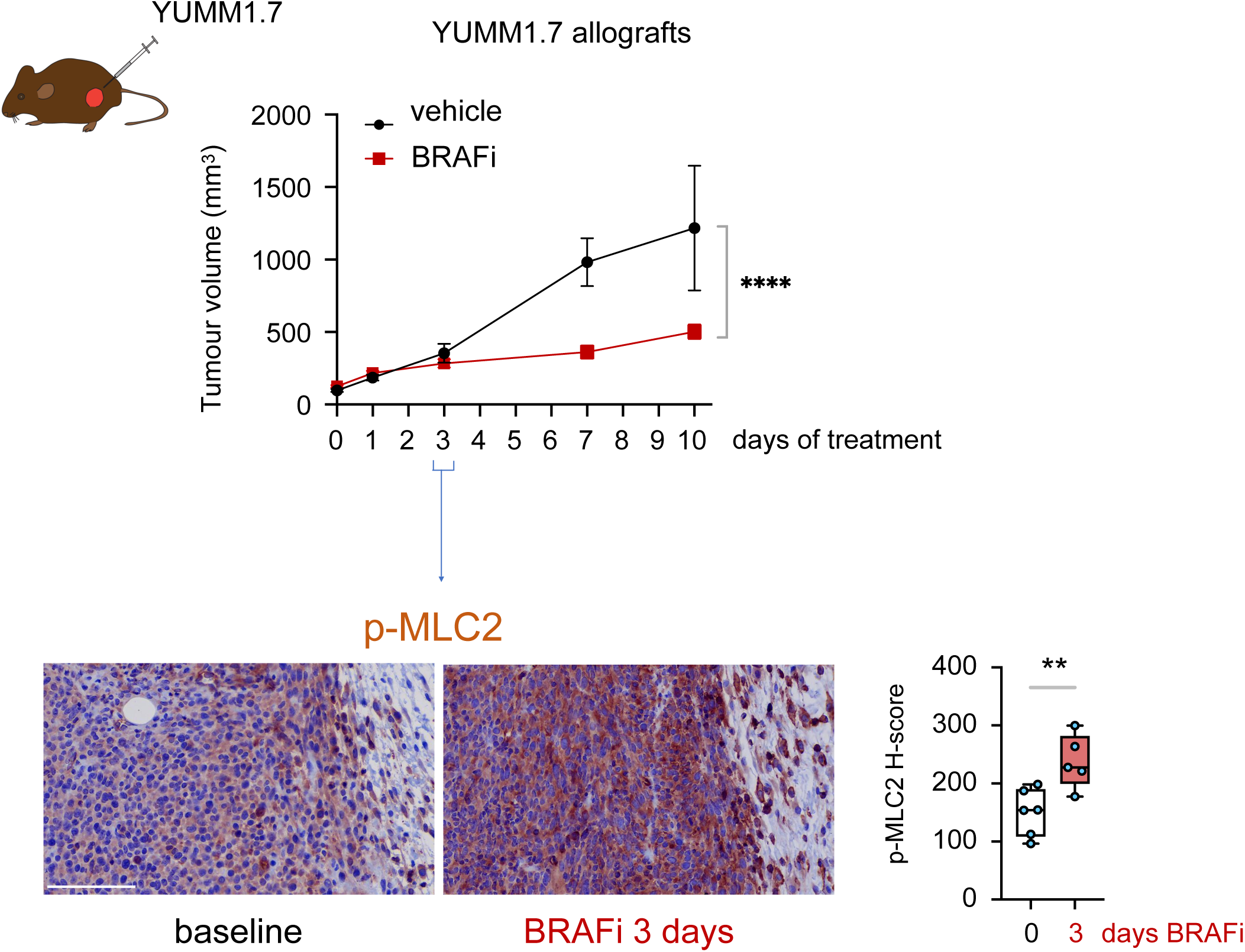
NMII levels increase early on during adaptation to MAPK inhibition in vivo. Top, graph shows tumour volume of YUMM1.7 allografts under BRAFi treatment. Bottom, representative images and quantification of p-MLC2 levels (H-score) by immunohistochemistry of YUMM1.7 allografts at baseline (day 0) and after 3 days of treatment. Boxplot: median (centre line); interquartile range (box); min-max (whiskers) and dots for individual data points (n= 6 tumours/group). P-value by two-tailed t-test. Scale bar, 100 μm.

Therefore, pp-MLC2 levels increase during the first 2 weeks of treatment, and this is generally followed by elevated MLC2 protein levels. The latter could be due to transcriptional changes triggered by elevated NMII activity, as shown in fibroblasts (43). We analysed expression levels of NMII genes in published transcriptomic data of melanoma cell lines during adaptation to BRAFi or BMi (14). Several NMII genes increased their expression upon 1-2 days BRAFi vs parental cells, and this was maintained in DTP/DTPP/resistant states (Fig. S2c). We confirmed by quantitative RT-PCR that MLC2 gene *MYL9* increased after 24 h BMi treatment in A375 cells, but this was prevented when cells were co-treated with transcription inhibitor actinomycin D (Fig. S2d), suggesting active transcription. We analysed published RNA-seq data of M397 melanoma cells treated with BRAFi for 72 h (44), which also included data of assay for transposase-accessible chromatin using sequencing (ATAC-seq), which provides information of accessible chromatin regions, usually associated with active transcription. *MYL9* mRNA was upregulated by BRAFi concurrently with an increased availability of a regulatory region in *MYL9* (Fig. S2e). Together, these data suggest that active transcription contributes, at least partially, to the induction of some NMII genes during adaptation to BMi.

We also assessed published transcriptomes of baseline and on-treatment human melanoma biopsies (14,45). Levels of several NMII genes increased in some BRAFi-treated vs baseline tumours (Fig. S3a). Analysis of published RNA-seq data of YUMM1.7 mouse melanoma allografts treated with BRAFi showed that MLC2 gene *MYL9* was upregulated in residual BRAFi-treated tumours vs controls (Fig. S3b).

Therefore, NMII activity increases early on during adaptation to BMi, due to changes in activity that are followed by transcriptional changes that may reinforce and maintain higher NMII activity. These data contribute to explain the increased NMII levels in BMR sub-lines (Fig. 1).

### NMII overactivation during adaptation to BMi is controlled by ROCK rather than by ERK rebound

Adaptation to MAPKi mostly involves an early reactivation of ERK1/2 activity (pp-ERK1/2) (6,46), although with varying degrees of pp-ERK1/2 recovery (14,47). In our BMi time courses, after an initial suppression, pp-ERK1/2 levels rebounded and followed similar kinetics to pp-MLC2 (Fig. 2), so we wondered if ERK rebound could contribute to overactivated NMII. We compared treatments of A375 cells with BMi or a triple therapy (BMEi) of BMi plus the clinically relevant ERK inhibitor (ERKi) SCH772984. BMEi reduced pp-ERK1/2 levels vs BMi alone (Fig. 4a). Nevertheless, levels of pp-MLC2 still increased under BMEi although to a lesser extent (Fig. 4a). Therefore, apart from a minor contribution of ERK, there may be additional mechanisms contributing to NMII activity dynamics during adaptation to BMi.

**Figure 4.**
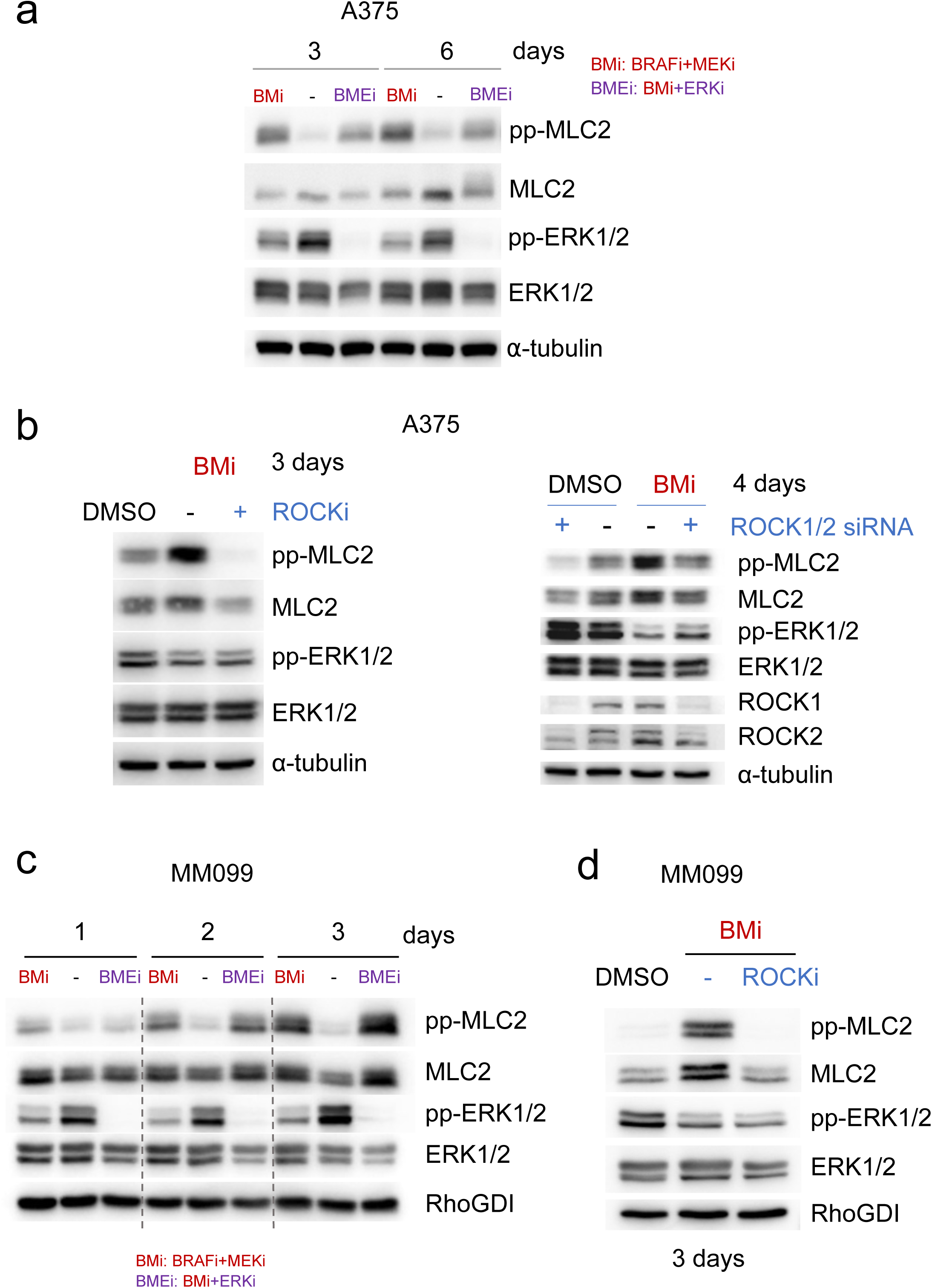
NMII overactivation during adaptation to BMi is controlled by ROCK rather than by ERK rebound. (**a, c**) Immunoblots of indicated proteins from melanoma cell lines after treatments with BMi (125 nM BRAFi dabrafenib + 6.25 nM MEKi trametinib) or BMEi (BMi plus 1 μM ERKi SCH772984). Representative of 3 independent experiments. (**b**) Immunoblots of indicated proteins from A375 cells treated with BMi for 2 days and additional 24 h in the presence or absence of 1 μM ROCKi GSK269962A (left panel, n=3); or treated with BMi for 3 days and transfected with siRNA against ROCK1 and ROCK2 (right panel). Non-targeting siRNA (-) was used as control. (**d**) Immunoblots of indicated proteins from MM099 cells treated with BMi for 2 days and additional 24 h in the presence or absence of 1 μM ROCKi GSK269962A. Representative of 3 independent experiments.

BMi treatment increased expression levels of several NMII regulators: MLCK in A375 cells, and MLCK and ROCK2 in several BMR vs P cells (Fig. S4a-b). Thus, we wondered if any of these kinases could be contributing to NMII dynamics during adaptation to BMi. Despite efficient MLCK protein knockdown, MLCK silencing in A375 cells via siRNA did not prevent the increase of pp-MLC2 levels upon BMi (Fig. S4c). However, ROCK1+2 inhibition (ROCKi) or silencing did block the induction of pp-MLC2 by BMi (Fig. 4b).

Then we tested a melanoma culture with dedifferentiated phenotype, which are usually intrinsically resistant to MAPKi (8,22,27,48). Consequently, a 24 h BMi treatment reduced pp-ERK1/2 levels to a much lesser extent in dedifferentiated MM099 cells than in A375 cells, but co-treatment with ERKi (BMEi) did decrease pp-ERK1/2 levels in MM099 cells (Fig. S4d). Nevertheless, in MM099 cells pp-MLC2 levels increased upon BMi treatment and were not affected when co-treated with ERKi (Fig. 4c). As in A375 cells, in MM099 cells ROCKi prevented NMII overactivation under BMi (Fig. 4d).

Furthermore, we assessed NMII gene levels in transcriptomic data of resistant lines classified in 2 groups: Rr (<u>R</u>esistant MAPK-redundant) that have weaker ERK activity rebound, higher RTK signalling and are more dedifferentiated vs parental cells; and Ra (<u>R</u>esistant MAPK-<u>a</u>ddicted) that have high ERK activity rebound and MAPK signalling, are more differentiated and more similar to parental cells (14). Generally, there were more NMII genes whose expression changed to a greater extent, especially upregulation, in Rr cells (less ERK signalling) than in Ra vs parental cells (Fig. S4e), further suggesting the independence of NMII dynamics and ERK activity during adaptation to BMi and also in the resistant state.

Therefore, during adaptation to BMi, the increase in NMII activity depends mostly on ROCK than on ERK or MLCK.

### NMII is not activated in melanomas with an hyperdifferentiated/pigmented phenotype during adaptation to BMi

During adaptation to BRAFi/BMi some melanoma cells switch to an hyperdifferentiated/pigmented state, which involves upregulation of MITF that promotes cell survival (12,49–51). A2058 melanoma cells promptly increased MITF protein levels under BMi and became pigmented, suggesting an hyperdifferentiated trajectory under BMi (Fig. 5a, S5a). Although ERK activity rebounded after 2 days of BMi treatment, pp-MLC2 were not recovered after 6 days of treatment and were still lower in A2058.BMR than in P cells (Fig. 5a-b, S5b). Interestingly, siRNA-mediated MITF knockdown in BMR cells increased pp-MLC2 levels (Fig. 5c), suggesting that elevated MITF during BMi-induced hyperdifferentiation prevents NMII overactivation.

**Figure 5.**
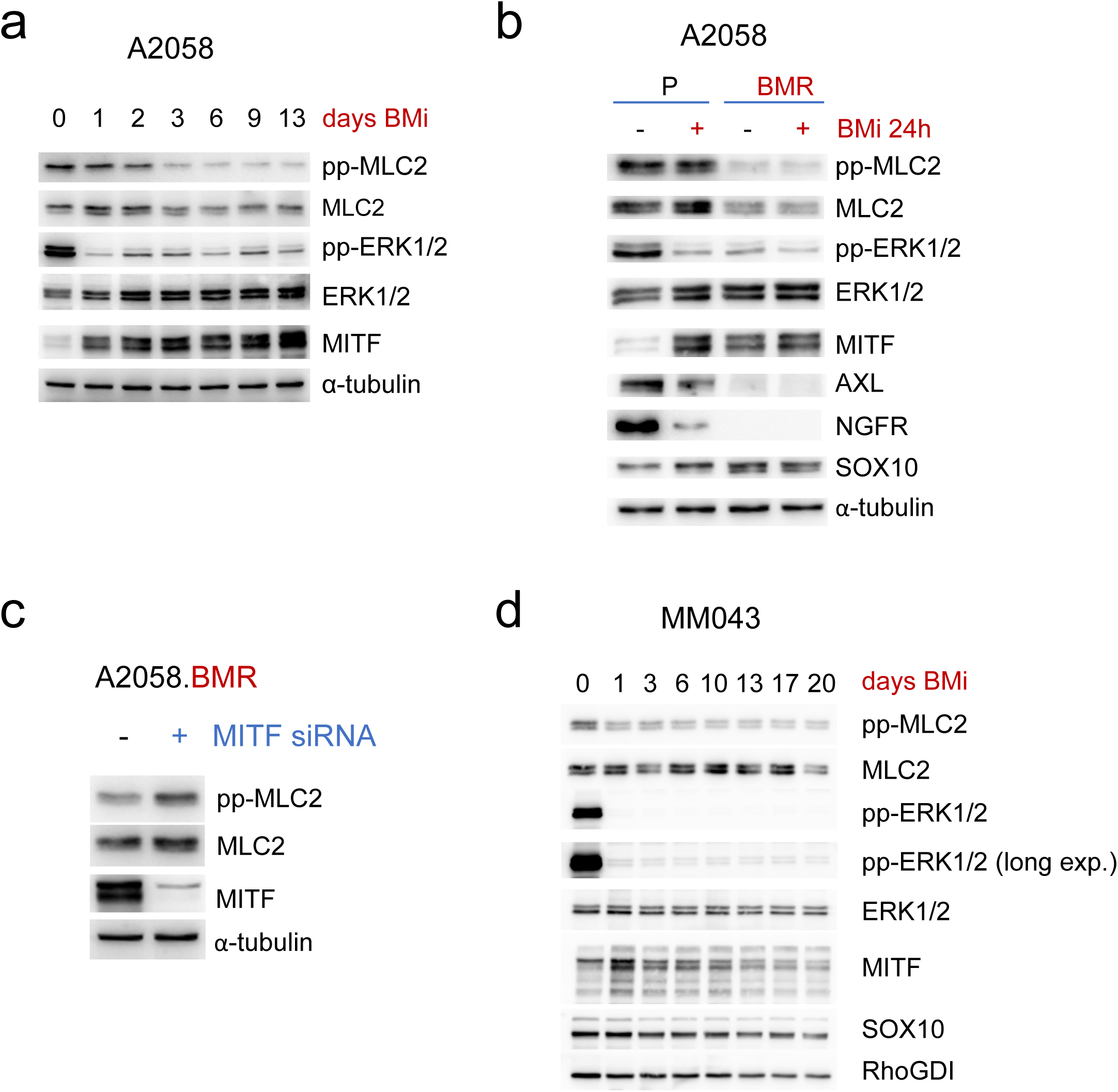
NMII is not activated in melanomas with an hyperdifferentiated/pigmented phenotype during adaptation to BMi. (**a-b, d**) Immunoblots of indicated proteins from melanoma cells after treatments with BMi (125 nM BRAFi dabrafenib + 6.25 nM MEKi trametinib). Long exp. = long exposure. (**c**) Immunoblotting of indicated proteins 72 h after transfection with MITF or non-targeting control siRNA (-). Representative of 3 independent experiments.

The melanocytic patient-derived culture MM043 (41) also followed an hyperdifferentiated/pigmented trajectory with elevated MITF protein levels, decreased pp-MLC2 and pp-ERK1/2 after 1 day of BMi treatment, and increased pigmentation after 3 days of BMi (Fig. 5d, S5c-d). However, pp-MLC2 remained low after 20 days of treatment, and cells were still pigmented.

Therefore, NMII activity is not increased in melanomas due to MITF-driven hyperdifferentiatiation during adaptation to BMi.

### NMII inhibition decreases tolerance of some melanomas to BMi

Since NMII activity increased very early on during adaptation to BMi in most models (8 out of 10), we hypothesised that co-targeting NMII activity along BMi could delay emergence of DTPs. Treatment of A375 cells with the widely used NMIIi blebbistatin along with standard of care BMi led to a lower percentage of surviving cells than under BMi alone for up to 21 days of treatment (Fig. 6a). In this model, blebbistatin mono-therapy was more efficient than BMi (Fig. S6a). Blebbistatin also reduced survival of A375 cells grown on collagen I matrices (Fig. S6b), which recapitulate the dermal environment in which melanomas grow and disseminate (52). Furthermore, siRNA against MLC2 also decreased cell survival (Fig. S6c), phenocopying the effects of blebbistatin.

**Figure 6.**
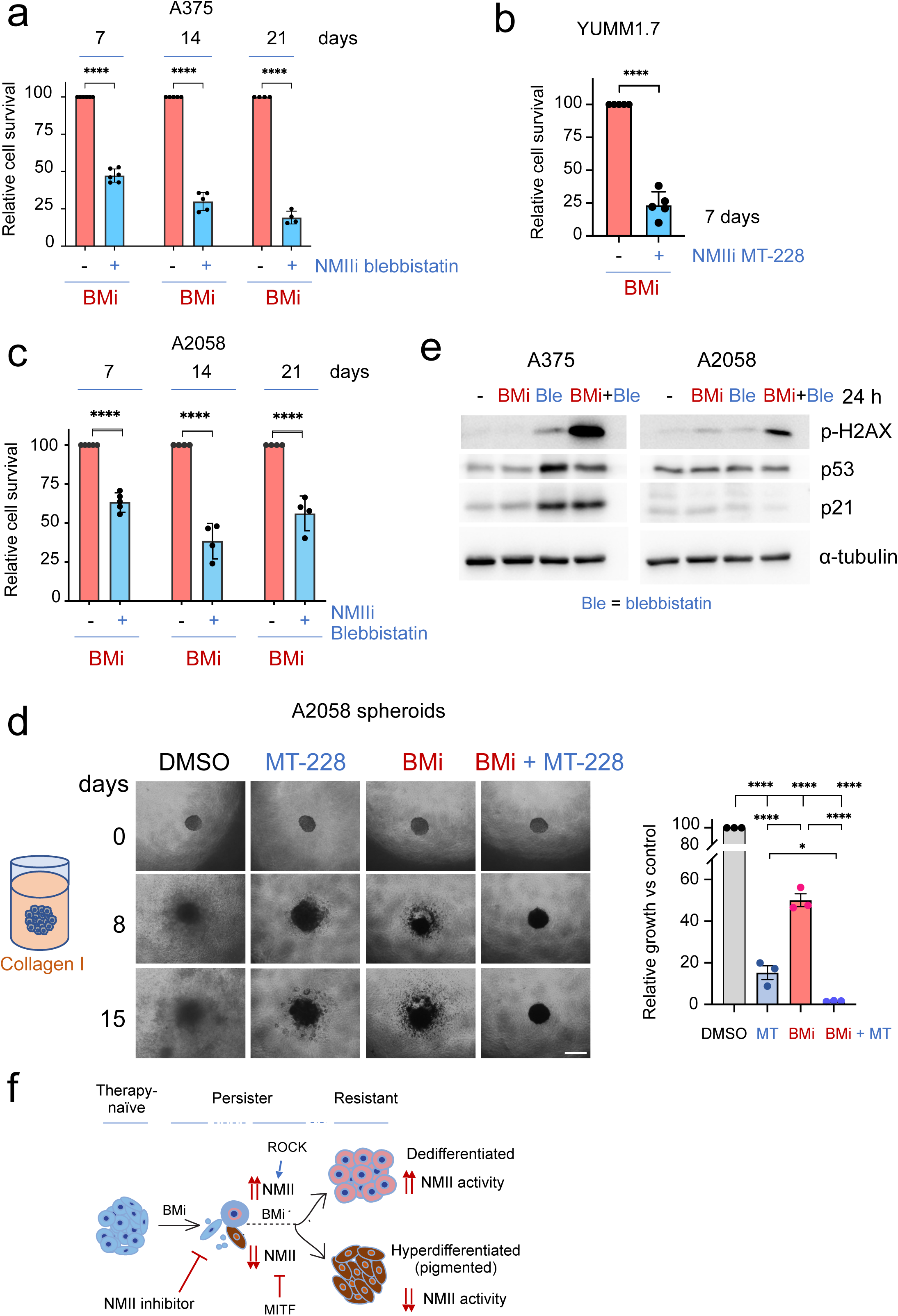
NMII inhibition decreases tolerance of some melanomas to BRAFi+MEKi. (**a, b, c**) Relative cell survival by crystal violet staining of melanoma cultures after treatment with BMi (125 nM BRAFi dabrafenib + 6.25 nM MEKi trametinib), 10 μM NMII inhibitor (blebbistatin or MT-228), or both (n=4-6). Dots indicate independent experiments. (**d**) Left, representative phase-contrast images of A2058 3D spheroids in collagen I matrix treated with MT-228, BMi or both. Scale bar, 400 μm. Right, quantification of spheroid growth (n=3). (**e**) Immunoblots of indicated proteins from melanoma cells after 24 h treatments with BMi, blebbistatin or both. Representative of 2 independent experiments. (**f**) Model summarising the main findings of this study. P-values by one-way ANOVA with Sidak (a, c) or Tukey post hoc test (d); two-tailed t-test (b). (*p<0.05, **p<0.01, ***p<0.001, ****p<0.0001).

We also tested a recently developed NMIIi, MT-228, which is more specific of non-muscle than cardiac myosin (53). Combination of MT-228 with BMi decreased survival more efficiently than BMi alone in several models (Fig. 6b and S6d).

To model potential pre-clinically relevant settings, we also tested sequential drug regimens in A375 cells with drugs switching every 3 or 7 days for up to 21 days, since alternating therapies may reduce chances of resistance to either drug (54,55). Initial treatment with blebbistatin and later BMi was more efficient than vice versa, and blebbistatin→BMi was as efficient as continuous treatment with the combination (Fig. S6e).

Surprisingly, combining blebbistatin with BMi was also more efficient than BMi alone in A2058 cells (Fig. 6c), which did not overactivate NMII under BMi (Fig. 5a). The combination BMi+MT-228 was also superior in reducing the growth of A2058 3D spheroids in collagen I compared to BMi alone (Fig. 6d).

Mechanistically, the combination blebbistatin+BMi for 24 h increased levels of p-H2AX, indicative of DNA damage (Fig. 6e). In A375 cells, this was concomitant with elevated p53 and p21 protein levels, suggesting again damage and cell cycle arrest. If unresolved, this damage could lead to cell death (56), which would explain the reduced long-term cell survival (Fig. 6a,c). A2058 cells harbour inactivating mutations in p53 (COSMIC COSS2651881), which may underlie the absence of p53 induction in this model.

Therefore, in some melanomas combining the standard of care BMi with a NMIIi delays the emergence of DTPs, and most likely, long-term acquired resistance.

## Discussion

Inhibition of the MAPK pathway generally leads to effective but short-lived responses due to development of resistance. Apart from pre-existing alterations, early adaptive, non-genetic mechanisms promote phenotype switching and cellular state transitions that allow survival of persister cells. Since persister/resistant cells undergo extensive cytoskeletal remodelling, new dependencies and vulnerabilities arise on these cells. Therefore, the cell cytoskeleton has emerged as a potential therapeutic target to overcome drug adaptation and resistance (24,35).

In this study, we find that NMII activity increases in BMR sub-lines that dedifferentiate during adaptation to BMi. NMII overactivation occurs relatively early during treatment, within the first 14 days, although in several models the increase occurs after 1-3 days of treatment. We also provide limited evidence that this occurs in vivo early during treatment, although this warrants confirmation in additional models and patient samples. A recent paper described NMII pathway overactivation and mesenchymal dedifferentiation in non-small cell lung cancer cells (NSCLC) during early adaptation to EGFR, KRAS or BRAF inhibition (57). Although this should be validated in other cancer models, our and this study suggest that NMII overactivation could be a general mechanism of adaptation to various targeted therapies. Further, NMII overactivation could be a biomarker of early adaptive resistance in cancer subtypes undergoing dedifferentiation. NMII activity does not increase in 2 models that adapt to BMi through hyperdifferentiation. Previous work showed overactivation of NMII in dedifferentiated melanoma cells, either BRAFi-resistant (36,38) or podoplanin-positive, therapy-naïve (58), which supports the overactivation of NMII in dedifferentiated BMR but not in hyperdifferentiated BMR cells.

Regarding possible links between NMII and phenotype switching, NMII activity increases before the modulation of phenotype markers in some models, but in others, levels of NMII and phenotype markers change concurrently. Therefore, we cannot conclude what occurs first, and further studies with additional models should clarify this question. Furthermore, parental UACC62 and SKMEL28 are both melanocytic but dynamics of NMII and phenotype markers differ substantially in these models, while dynamics are similar in A375 (NCSC) and UACC62. Although additional models will be required, the baseline phenotype does not seem to be linked to dynamics of NMII and phenotype switching under BMi.

The increase of NMII during adaptation to BMi seems to occur largely independently of ERK rebound, as shown in the ERKi experiments. Further, in the A2058 model ERK rebounds but NMII activity decreases, and expression of NMII genes increase to a larger extent in resistant lines that have weaker ERK signalling reactivation (MAPK-redundant). Perhaps high NMII activity is not necessary in the cases with enough MAPK reactivation (MAPK-addicted), and/or in models in which there is a bypass of ERK pathway through activation of downstream pro-survival pathways such as MITF (8). Our study suggests that MITF prevents NMII activation under BMi treatment, in line with a previous paper showing that MITF depletion in parental melanomas triggers ROCK-dependent invasion (59), although this study did not assess NMII activity.

MLCK could also be contributing to NMII overactivation since MLCK levels increase in several models during adaptation/resistance to BMi. However, MLCK silencing does not prevent NMII overactivation. This could be due to other kinases known to regulate NMII activity still being present, such as ROCK (32), whose blockade does prevent BMi-induced NMII overactivation. Since ROCK regulates NMII activity in several parental (60–63) and BRAFi-resistant melanoma sub-lines (39), perhaps ROCK control on NMII dominates over other potential regulators. Consistent with this, ROCKi impairs EGFRi-induced NMII activation and actin fibre formation in NSCLC DTPs (57).

Since ROCKi prevents NMI overactivation under BMi, co-targeting ROCK with BMi could delay resistance to BMi, as shown for BRAFi and ERKi (64). ROCK activity could be promoted by upstream Rho GTPases, as some Rho GTPases are overactivated/overexpressed in DTPs (57,65), BRAFi-resistant (29,66) and BMR (47) melanomas. However, ROCK function could still be compensated by other parallel cytoskeletal regulators that contribute to MAPKi resistance, such as PAK (47) or FAK (10,65,67). Therefore, we propose a direct NMII inhibition using novel and non-muscle specific NMIIi such as MT-228 (53) or MT-125 (68). MT-125 has shown promising anti-tumour effects in glioblastoma mouse models and is now under clinical evaluation (NCT07185880).

Mechanistically, there is elevated DNA damage after combined NMIIi+BMi. This damage could be caused by increased reactive oxygen species after NMII inhibition (68,69), which, concurrently with suppression of DNA repair pathways upon MAPKi (70), would explain the reduced survival with NMIIi+BMi.

Although further in vivo validation will be needed, our study suggests that melanomas with low/moderate sensitivity to BMi may benefit from co-targeting NMII. Alternating NMIIi→BMi schedules are as efficient as continuous NMIIi+BMi treatment. However, intermittent dosing should also be tested in case this delays development of resistance to either drug, at least as shown in preclinical settings for BMi (54,55).

Co-targeting NMII with BMi is effective even in melanomas that hyperdifferentiate and do not overactivate NMII, highlighting the need to assess the dependence on intrinsic sensitivity to NMIIi. In fact, we observe a range of responses after NMIIi monotherapy: while survival of A375 cells is greatly compromised, A2058 cells are moderately affected and the survival of SKMEL28 or WM88 cells is unaffected (Fig. S6f). The greatest responses seem to be in the most dedifferentiated phenotypes (mesenchymal and NCSC), although further analyses using a larger panel of melanomas should be performed to strengthen this statement. Therefore, and to reformulate the conclusion above, our study suggests that melanomas with low/moderate sensitivity to BMi and medium/high sensitivity to NMIIi would most likely benefit from the combination NMIIi+BMi.

In summary, we identify NMII overactivation as a marker of early adaptation to MAPK inhibition in dedifferentiating melanomas. In some melanomas, combined BMi+NMIIi delays emergence of persister cells (Fig. 6f). These findings pave the way for future proof-of-concept studies using combinatorial approaches to delay or prevent the development of resistance to MAPK-targeted therapy.

## Acknowledgments

JLO acknowledges the following funding: Atracción de Talento Investigador fellowship (2019-T1/BMD-13642 and 2023-5A-BMD-28922) by Comunidad de Madrid; Grant PID2021-122306OB-I00 funded by MICIU/AEI/10.13039/501100011033 and by FEDER, EU; Grant CNS2023-143636 funded by MICIU/AEI/10.13039/501100011033 and by European Union NextGenerationEU/PRTR. MDR was supported by “Programa INVESTIGO. Mecanismo de Recuperacion y Resilencia. Comunidad de Madrid” (Project #39) funded by European Union NextGenerationEU/PRTR. AM and RML were funded by Instituto de Salud Carlos III (ISCIII) through the projects “PI18/00573; PI21/00294; PI24/1433” (Co-funded by European Regional Development Fund/European Social Fund “A way to make Europe”/” Investing in your future”) and co-funded by the European Union. OM was supported by Barts Charity (MGU0504). We are grateful to Victoria Sanz-Moreno, Ghanem Ghanem, Marisol Soengas, Sophia Karagiannis, Richard Marais, Oystein Fodstad and Mary J Hendrix for providing cell lines; Luis del Peso for advice and his PCR thermocycler; Bruno Sainz Jr and Diego Navarro for helping with slide scanning; Victoria Sanz-Moreno, Anna Obenauf, Oskar Marin-Bejar, Miguel Vicente-Manzanares and Barbara Acosta-Iborra for advice and/or helpful discussions; and the facilities at IIBM Sols-Morreale for technical support.

## Author contributions

AGP carried out most of the experiments and generated some BMR sub-lines; MDR, LSG, SNA, ADL carried out experiments with ERKi; LSG conducted 3D spheroid assays; EJY, LR, CAM synthesised MT-228; MCS analysed ATAC-seq data; RMM, AM, JG, CR performed in vivo experiments; EPG, CDU, FS, JMMG, MPL conducted immunohistochemistry on tumour sections that were analysed by OM; JLO generated most BMR sub-lines, carried out initial time-course and cell survival experiments, and analysed transcriptomic data; JLO conceived, coordinated and supervised the whole study, obtained main funding, and wrote the manuscript and drafted figures with the help of AGP; all authors read and approved the manuscript.

## Competing Interests

EJY and CAM hold equity in Myosin Therapeutics, which holds the license to MT-228. No other authors declare known competing interests.

## Methods

Supplementary Tables include details and sources for cell lines (Table S1), small molecular inhibitors and other reagents (Table S2), siRNA (Table S3), antibodies (Table S4) and gene expression data (Table S5).

### Cell culture

Commercially available melanoma cell lines were acquired from other research groups (Table S1) and authenticated upon expansion by short-tandem repeat (STR) genotyping with the StemElite ID kit (Promega) by the IIBM Genomics Facility. STR genotyping of mouse YUMM1.7 cells was performed by Microsynth (Switzerland). BRAFi PLX4720-resistant sub-lines were previously characterized (39).

Short-term patient-derived melanoma cultures (MMxxx) have been extensively characterised (40,41,71). Some MMxxx were provided by Prof. Dr. Ghanem Ghanem via a material transfer agreement and others were acquired through CancerTools UK. These cultures had been derived from patient biopsies by the Laboratory of Oncology and Experimental Surgery (Institute Jules Bordet, Brussels).

MMxxx cultures were cultured in low tyrosine media (Ham’s F10) to prevent phenotype drift (72). The other cell lines were grown in DMEM. Both media were supplemented with 10% fetal bovine serum (FBS) and penicillin+streptomycin (“complete medium”), and cells were grown in an incubator at 37° C, 5% CO2 and 95% humidity. All cultures were confirmed to be mycoplasma-free at least once a month.

### Small molecule inhibitors

The following small molecule inhibitors were used in in vitro experiments: BRAFi dabrafenib (GSK2118436), encorafenib (LGX818) and PLX4720; MEKi trametinib (GSK1120212) and binimetinib (MEK162); ERKi SCH772984, NMIIi (-)-blebbistatin, ROCKi GSK269962A. MT-228 was synthesized in-house as described (53). Drugs were resuspended in pure DMSO at 10 mM (except NMIIi at 50-100 mM), and these stocks were kept frozen in polypropylene tubes at -80° C. Blebbistatin and MT-228 were kept in polypropylene amber tubes and wrapped in parafilm to prevent moisture. Stocks were diluted in fresh media right before addition to the cells.

### Generation of BRAFi+MEKi-resistant sub-lines

BRAFi+MEKi-resistant sub-lines (BMR) were generated after chronic exposure of parental lines to 125 nM dabrafenib and 6.25 nM trametinib (except WM983A and WM88, 1 μM dabrafenib + 50 nM trametinib). Cells were seeded in T25 flasks, treated the following day with fresh drugs, and medium was replenished every 2-3 days. Every 3-4 weeks cells were carefully detached and seeded onto new flasks, to prevent contaminations from being in the same flask for too long. After 1-5 months, cells gradually were able to grow under BMi, allowing generation of frozen stocks. Then, BMR cells were confirmed to have less sensitivity to BMi compared to their parental counterparts. BMR were continuously grown in the presence of BMi unless otherwise stated. Most BMR sub-lines grew faster in the absence of BMi (Fig. S1a), as previously reported (55,73,74). BMR sub-lines were authenticated by STR genotyping at the time of characterisation (Table S1).

### Cell culture on collagen matrices

Cells were grown on collagen I matrices as described (39,61). Briefly, bovine collagen I thick gels were polymerized at 1.7 mg/mL (for 1 mL of gel, 0.55 mL of collagen stock were mixed with 0.2 mL 5x DMEM, 0.215 mL milli-Q water, 0.03 mL 0.1N NaOH). This solution was de-gased using a line vacuum, dispensed onto culture plates (0.3 mL/T24-well or 0.1 mL/T96-well) and incubated at 37° C for 3-4 h. Cells were seeded on top of gels at 10,000 cell/well and drug treatments started 16 h later. Calculations of final drug concentrations took into account the volume of collagen. At endpoint, collagen I gels were washed with PBS and fixed with 4% formaldehyde for 15 min, and phase-contrast images were taken.

### 3D spheroids in collagen matrices

Spheroid culture was carried out by the hanging drop method. A2058 cells were resuspended in low viscosity medium at 25,000/mL. Low viscosity medium was made by mixing 800 μL of complete medium plus 200 μL of low viscosity methylcellulose solution (1.2 g methylcellulose/100 mL DMEM). Then, 20 μL droplets (500 cells) were suspended from the lid of tissue culture dishes for 72 h at 37° C, 5% CO_2_, allowing cell clustering into compact spheres. PBS was added to the dishes to limit evaporation. Spheroids were collected and added to 100 μL of a collagen I solution (1.7 mg/mL) in 96-well plates and left in the incubator at 37° C for 4 h. Afterwards, 100 μL of medium with 10% FBS and drugs were added on top of the gel and phase contrast pictures were taken every 2-3 days. For quantification of spheroid invasion/growth, the increase on the area occupied by the spheroid between day 0 (when spheroids were embedded into the collagen I matrix) and indicated time point was calculated by using ImageJ software.

### Cell survival analysis by crystal violet staining

Long-term survival was performed on tissue culture plastic dishes unless otherwise specified. Cells were seeded in multi-well plates (1,000-2,000 cells/cm^2^) and treated for several days/weeks, re-adding drugs in fresh media every 2-3 days. Then cells were washed with PBS, fixed with 4% formaldehyde for 15 min, washed twice and stained with 0.05% crystal violet for 15 min. Plates were rinsed thoroughly with tap water and air-dried overnight. Crystal violet was solubilized using 10% acetic acid, and absorbance at 595 nm was quantified using a SPECTROstar Nano plate reader (BMG LABTECH).

### RNAi transfection

Cells were seeded (100,000 cells/6-well) and were transfected the next day with 20 nM siRNA oligonucleotides and 7.5 μL RNAiMax in Opti-MEM Reduced Serum Medium (Gibco). After 48-72 h cells were lysed to assess knockdown efficiency. For long-term survival (6 days), cells were transfected 3 days after the first transfection, and plates were fixed and stained with crystal violet as described above. ON-TARGETplus SmartPool siRNA and the non-targeting siRNA #1 were from Horizon Discovery (Table S3). Regarding experiments with siRNA under BMi treatment, cells were treated with BMi, next day siRNA were transfected, and 2 days later cells were lysed for immunoblotting.

### Antibodies

Antibodies, source and dilutions are described in Table S4.

### Immunofluorescence and confocal imaging

Cells were seeded on glass coverslips and treated as indicated in figure legends. At endpoint, coverslips were washed with PBS and were fixed with 4% methanol-free formaldehyde for 15 min and washed 3 time with PBS. Cells were permeabilised with 0.2% Triton X-100 for 5 min, blocked with 5% BSA-PBS for 1 h at room temperature (RT), and incubated with primary antibodies diluted in 5% BSA-PBS overnight at 4° C in wet chambers. Coverslips were washed and incubated with Alexa-488 or -647 anti-rabbit secondary antibodies for 1 h at RT. F-actin was detected with 546-Phalloidin (1 h RT) and nuclei were stained with DAPI. Coverslips were mounted in Prolong Diamond Antifade. Imaging was carried out on a Zeiss LSM 710 confocal microscope (Carl Zeiss) with Plan-Apochromat 40x/1.3 NA (Oil) dic objective lens and Zen software (CarlZeiss).

### Immunoblotting

Cells were washed once with cold PBS and lysed in Laemmli buffer (60 mM Tris pH 6.8, 2% SDS, 10 % glycerol). Plates were scraped and lysates were collected and frozen at -80° C. Lysates were then boiled, sonicated for 30 s (Bioruptor diagenode UCD-200, high setting) and protein concentration was quantified with DC Protein Assay Kit II. At least 20 μg of cell lysates in sample buffer with 100 mM DTT were fractionated in SDS-PAGE gels and subsequently wet-transferred to Immobilon-P PVDF filters. Membranes were blocked in 5% BSA in wash buffer (0.1% Tween 20-TBS). Primary antibodies were incubated overnight at 4° C. Membranes were washed with wash buffer and incubated with horse peroxidase-conjugated secondary antibodies for 1 h at RT. For detection, membranes were incubated with Clarity ECL and imaged using Vilber Fusion Solo-6S imager. Blots were stripped using a mild stripping buffer (50 mM Glycine, 2% SDS, pH 2) for 15 min, then washed extensively, and incubated with further primary antibodies with a different molecular weight than the previous one. Bands were quantified using ImageJ. Levels of pp-MLC2 and MLC2 were calculated after correction to the loading control.

### Quantitative RT-PCR

RNA was isolated using TriZol and treated with DNase Treatment and Removal Reagent following manufacturer’s instructions. One μg RNA was reverse-transcribed to cDNA using RT NZY First-Strand cDNA Synthesis Kit (NZYTech # MB12501) and diluted 1:10. Quantitative RT-PCR was carried out in a final volume of 10 μL/well in MicroAmp Fast Optical 48-Well Reaction Plates, including 12 ng cDNA as template, Power SYBR Green PCR Master Mix and 400 nM primers (*MYL9* and BM2 (Table S2)). PCR amplifications were carried out in a StepOne Real-time PCR System (Applied Biosystems). Data were analysed with StepOne software and expression levels were calculated using ΔΔCT, using *B2M* as reference.

### Mouse allografts

Animal experimentation in this study complies with all relevant ethical regulations, and was carried out in accordance with the principles of the Basel Declaration and recommendations of the Catalan Government (Generalitat de Catalunya, 1201/2005) concerning the protection of animals for experimentation. C57BL/6 mice were maintained by brother-sister mating under specific pathogen-free conditions at the University of Lleida (Spain). The protocol was approved by the Committee on the Ethics of Research in Animal Experimentation of the University of Lleida. One million YUMM1.7 cells were injected by subcutaneous injections into each flank of C57BL/6 mice (Charles River; half males and half females). Mice were monitored daily and when tumours were palpable, length and width were measured with a caliper to calculate tumour volume (mm3) = (length x width) x0.5. Seven days post-injection, tumours were on average 100 mm^3^ and were allocated into vehicle and 50 mg/kg vemurafenib (PLX4032) treatment arms. At indicated time points, mice were sacrificed and the tumours were collected and fixed in formalin and embedded in paraffin for subsequent immunohistochemistry.

### Immunohistochemistry on tumour allografts

Formalin-fixed, paraffin-embedded 5 μm sections were stained using standard immunohistochemistry methods. Antigen retrieval was performed in a pressure cooker using Tris-EDTA Buffer (10 mM Tris Base, 1mM EDTA, 0.05% Tween 20, pH 9.0). Blocking buffer (1% BSA, TBS 1x) and BLOXALL blocking solution were used to block nonspecific proteins and endogenous peroxidase and alkaline phosphatase activities. Diluted primary antibodies (p-MLC2) were incubated overnight at 4° C. Antibody detection was performed using ImmPRESS HRP Horse Anti-Rabbit IgG Polymer Detection Kit and ImmPACT NovaRED Substrate Kit, Peroxidase (HRP). Slides were counterstained with Mayer’s hemalum solution and mounted with Permount mounting medium. Then slides were scanned at 40x with a Motic EasyScan One slide scanner.

Quantification of p-MLC2 staining was carried out using QuPath v0.6.3 (75). Images were analysed by performing positive cell detection, and three different thresholds were applied according to the intensity scores (0, 1, 2 and 3). Next, the software was trained by creating random trees classification algorithm combined with the intensity information, in order to differentiate tumour from stroma and immune cells. Values used in the analysis correspond to the quantification of p-MLC2 in the invasive front.

### Gene expression analyses

Normalised RNA-seq data were downloaded from Gene Expression Omnibus (GEO), accession numbers GSE75313 (14); GSE127988 (76); GSE64741 (77); GSE134459 (44); and the European Genome-phenome Archive (EGA) accession number S00001000992 (45). More details are in Table S5. To overcome noise in differential expression values caused by extremely low levels, we added a pseudo-count value of 0.1 to all expression values, according to the protocol in (14). Then relative expression data was calculated as log_2_ fold change of treatment/resistant condition vs baseline/parental condition. Heapmaps were generated in Morpheus https://software.broadinstitute.org/morpheus/.

ATAC-seq data of M397 cells treated with BRAFi were downloaded from GSE134459 (44) and processed as follows. Raw read quality was assessed with FastQC. Adapter trimming was performed using Trim Galore v0.6.10 (cutadapt wrapper). Only successfully trimmed read pairs were used. Trimmed paired-end reads were aligned to the hg38 reference genome using Bowtie2 v2.5.2 with the very-sensitive preset and a maximum fragment length of 2000 bp. Alignments were piped to samtools for coordinate sorting, and alignment metrics were obtained using flagstat and idxstats. Aligned BAMs were deduplicated and filtered using samtools v1.19.2, retaining high-quality reads with MAPQ ≥ 30. Accessible chromatin regions were identified using MACS3 with ATAC-seq-appropriate parameters, including paired-end mode and q<0.01 for peak calling, generating narrowPeak, summits, and XLS files for each sample. Normalised genome-wide signal tracks were produced using bamCoverage (deepTools) with CPM normalisation, 10-bp binning, duplicate exclusion, and fragment-length extension. For locus-level visualisation, narrowPeak intervals and CPM-normalised bigWig tracks were imported into RStudio and plotted using standard genomic visualisation packages, enabling comparison of peak intensity across samples.

### Quantification and statistical analysis

GraphPad Prism (GraphPad Software) was used to carry out statistical analyses. Details of statistical analysis performed are in the figure legends. Bar graphs report mean ± SEM with individual data points as explained in figure legends. In figure legends, ‘‘n’’ means number of independent experiments unless otherwise stated.

### Data availability

The datasets used and/or analysed during the current study are available from the corresponding author on reasonable request.

