## Supplementary Figures for "Early overactivation of non-muscle myosin II during adaptation to combined BRAF and MEK inhibitors in dedifferentiating cutaneous melanomas"

### Figure S1

a

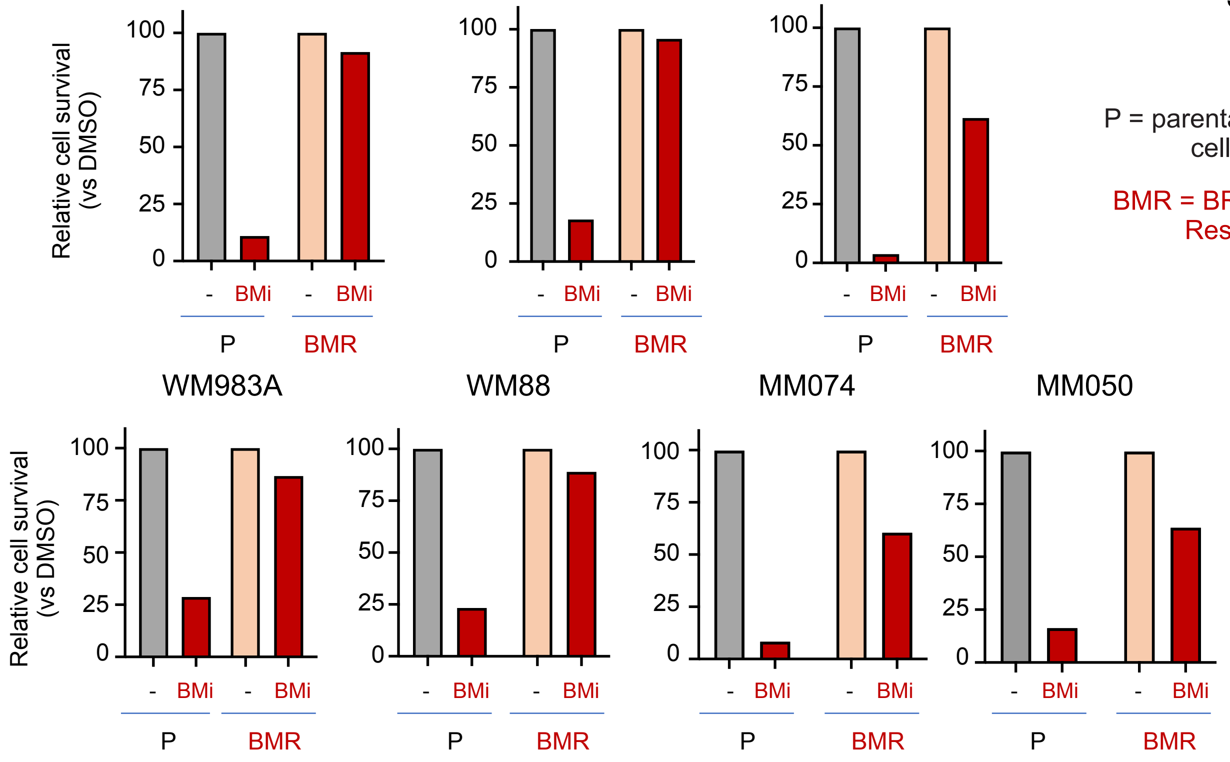


P = parental melanoma cell lines

BMR = BRAFi+MEKi-Resistant

b

| Phenotype | MITF | SOX10 | NGFR | AXL |
|---|---|---|---|---|
| **Melanocytic** | HIGH | HIGH | LOW | LOW |
| **Intermediate/Transitory** | HIGH | HIGH | HIGH | LOW |
| **Neural Crest Stem Cell NCSC** | LOW | HIGH | HIGH | HIGH |
| **Dedifferentiated Mesenchymal** | LOW | LOW | LOW | HIGH |

Differentiation

Resistance to BRAFi

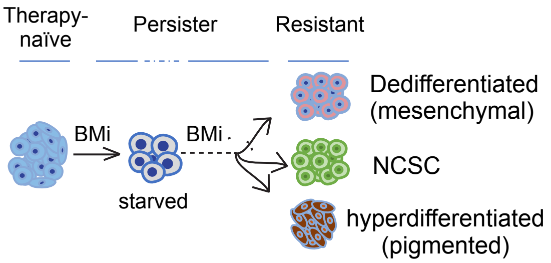


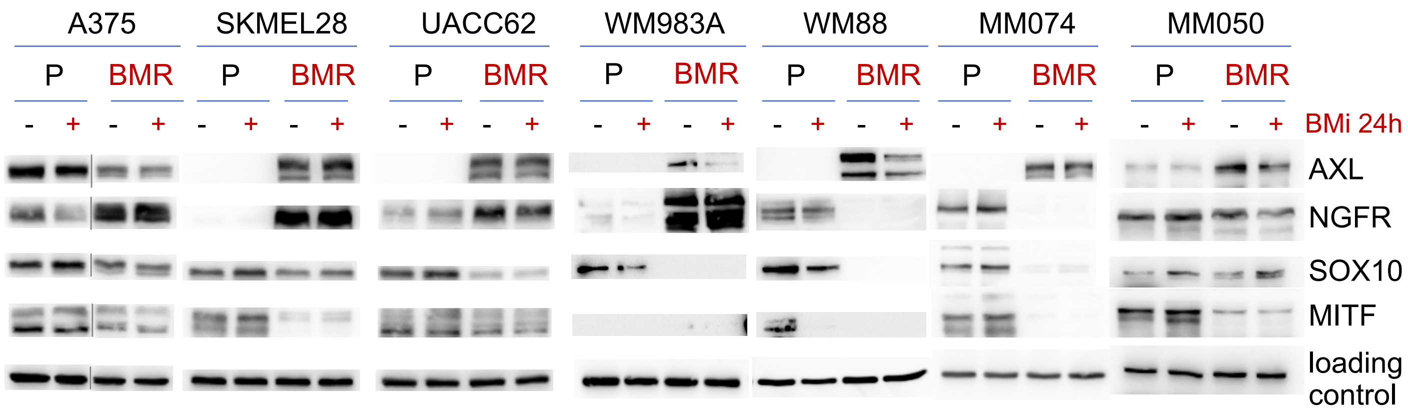


c

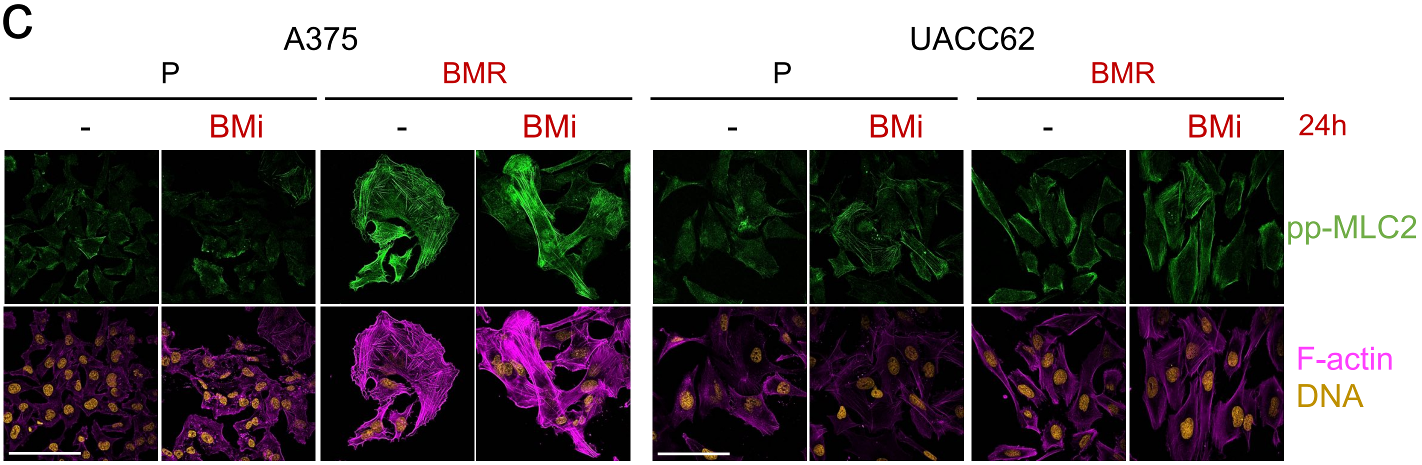

**Figure S1. BRAFi+MEKi-resistant melanoma sub-lines have higher levels of NMII activity than their parental counterparts.** (**a**) Relative cell survival by crystal violet staining of parental (P) and BMR cultures after BMi treatment for 3 days (except UACC62, 7 days) compared to vehicle control (DMSO, -). Representative of 2 independent experiments. (**b**) Top left, schematic of levels of phenotype markers in each phenotype (1–3). Top right, schematic of phenotype switching during adaptation to BMi (adapted from (4)). Bottom, immunoblots of indicated phenotype markers from pairs of parental (P) and BMR lines after 24 h treatment with BMi (125 nM BRAFi dabrafenib + 6.25 nM MEKi trametinib). Loading control was $\alpha$-tubulin except for WM983A and MM074 that had RhoGDI. Some of these (SKMEL28, WM983A, WM88, MM074) show the same loading control as main Fig. 1a since the phenotype markers were reprobed on the same blots. Lines in A375 blots denote non-contiguous lanes in the same gel/blot. (**c**) Confocal immunofluorescence images of pp-MLC2, F-actin (phalloidin) and DNA (DAPI) stainings after 24 h treatment with BMi. Scale bars, 100 $\mu$m.

### Figure S2

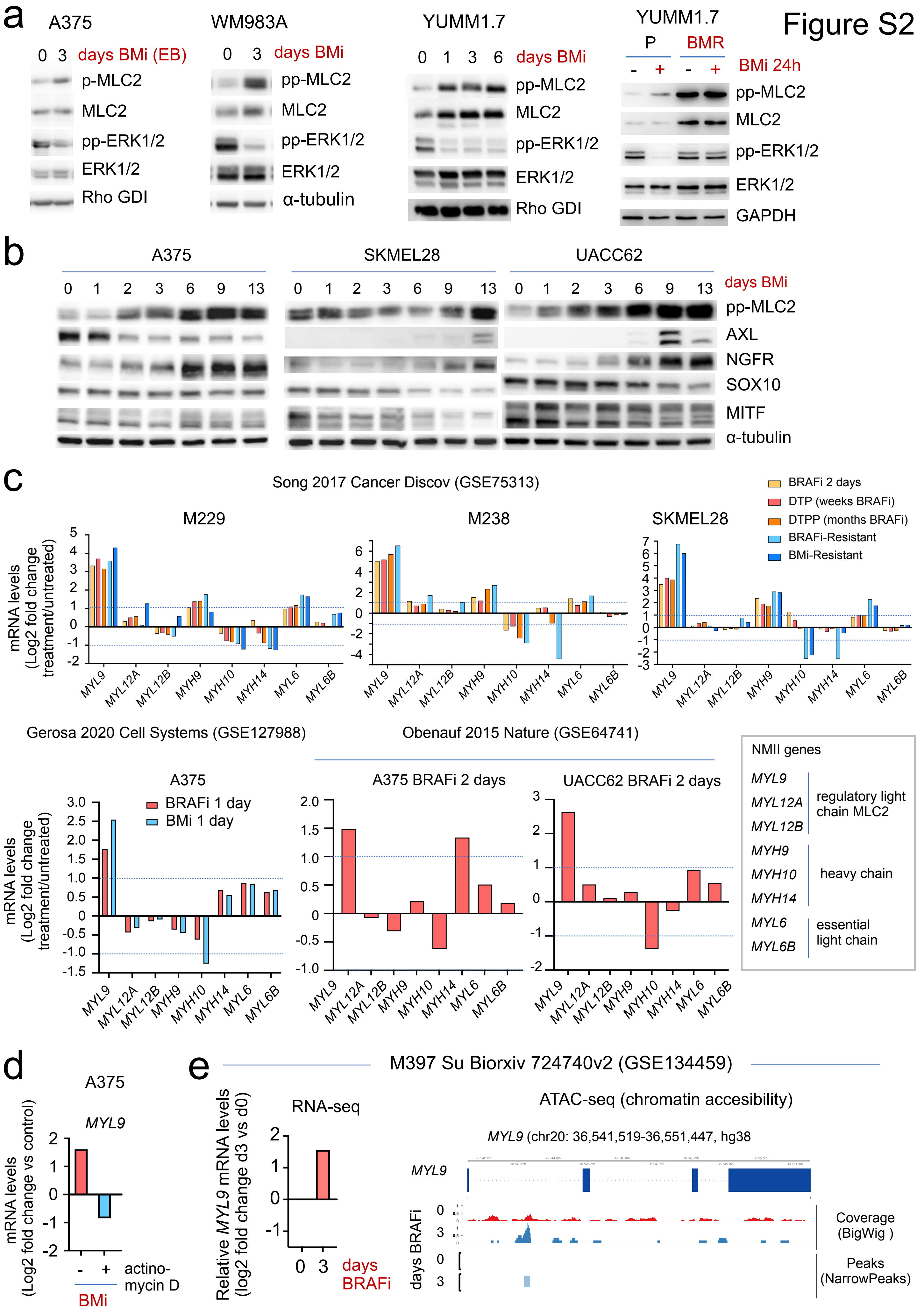
a
A375
0 3 days BMi (EB)
p-MLC2
MLC2
pp-ERK1/2
ERK1/2
Rho GDI
WM983A
0 3 days BMi
pp-MLC2
MLC2
pp-ERK1/2
ERK1/2
α-tubulin
YUMM1.7
0 1 3 6 days BMi
pp-MLC2
MLC2
pp-ERK1/2
ERK1/2
Rho GDI
YUMM1.7
P BMR
- + - + BMi 24h
pp-MLC2
MLC2
pp-ERK1/2
ERK1/2
GAPDH
b
A375
SKMEL28
UACC62
0 1 2 3 6 9 13 days BMi
pp-MLC2
AXL
NGFR
SOX10
MITF
α-tubulin
c
Song 2017 Cancer Discov (GSE75313)
BRAFi 2 days
DTP (weeks BRAFi)
DTPP (months BRAFi)
BRAFi-Resistant
BMi-Resistant
M229
M238
SKMEL28
mRNA levels (Log2 fold change treatment/untreated)
MYL9 MYL12A MYL12B MYH9 MYH10 MYH14 MYL6 MYL6B
Gerosa 2020 Cell Systems (GSE127988)
Obenauf 2015 Nature (GSE64741)
A375
BRAFi 1 day
BMi 1 day
A375 BRAFi 2 days
UACC62 BRAFi 2 days
NMII genes
MYL9
MYL12A
MYL12B
regulatory light chain MLC2
MYH9
MYH10
MYH14
heavy chain
MYL6
MYL6B
essential light chain
d
A375
MYL9
mRNA levels (Log2 fold change vs control)
- + actino-mycin D
BMi
e
M397 Su Biorxiv 724740v2 (GSE134459)
RNA-seq
Relative MYL9 mRNA levels (log2 fold change d3 vs d0)
0 3 days BRAFi
ATAC-seq (chromatin accesibility)
MYL9 (chr20: 36,541,519-36,551,447, hg38)
MYL9
days BRAFi
0
3
Coverage (BigWig )
Peaks (NarrowPeaks)

**Figure S2. NMII levels increase early on during adaptation to BRAFi+MEKi.** (**a-b**) Immunoblots of indicated proteins from parental melanoma cell lines after treatments with BMi (125 nM BRAFi dabrafenib + 6.25 nM MEKi trametinib, except A375 EB (1 μM Encorafenib + 0.3 μM Binimetinib)). pp-MLC2 blots in b are the same as in Fig. 2a, these are included here for comparison of kinetics vs phenotype markers. (**c**) Expression levels of NMII genes in published RNA-seq data of melanoma cultures after treatment with BRAFi for different times (1-2 days, weeks-months (drug-tolerant persister (DTP), drug-tolerant proliferating persister (DTPP))) and of BRAFi- and BMi-resistant sub-lines (5–7). Levels are plotted as $\log_2$ fold change of treated or resistant cells compared to parental or vehicle-treated cells. Dashed lines indicate changes larger or equal than two-fold. (**d**) Expression levels of MLC2 gene *MYL9* by quantitative RT-PCR from A375 cells treated for 15 h with BMi (1 μM + 50 nM) (or DMSO as control) and with or without inhibitor of transcription actinomycin D (5 μg/ml). mRNA levels are plotted as $\log_2$ fold change of treated vs double vehicle-treated cells (DMSO and no actinomycin). (**e**) Expression levels (RNA-seq) and chromatin availability (ATAC-seq) of *MYL9* from published data (8). Expression levels are plotted as $\log_2$ fold change of BRAFi day 3 vs day 0. ATAC-seq analysis shows accessible chromatin region within *MYL9* in BRAFi day 3.

Figure S3

a

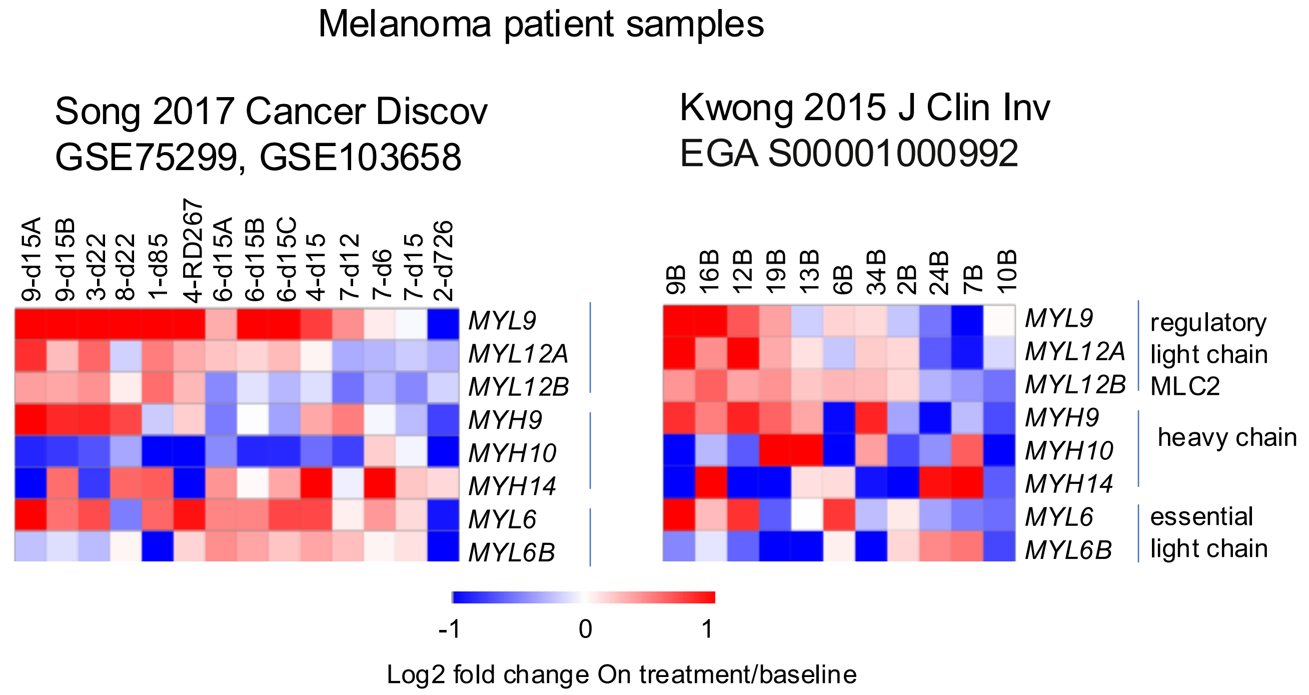
Melanoma patient samples
Song 2017 Cancer Discov
GSE75299, GSE103658
9-d15A
9-d15B
3-d22
8-d22
1-d85
4-RD267
6-d15A
6-d15B
6-d15C
4-d15
7-d12
7-d6
7-d15
2-d726
MYL9
MYL12A
MYL12B
MYH9
MYH10
MYH14
MYL6
MYL6B
Kwong 2015 J Clin Inv
EGA S00001000992
9B
16B
12B
19B
13B
6B
34B
2B
24B
7B
10B
regulatory
light chain
MLC2
heavy chain
essential
light chain
-1
0
1
Log2 fold change On treatment/baseline

b

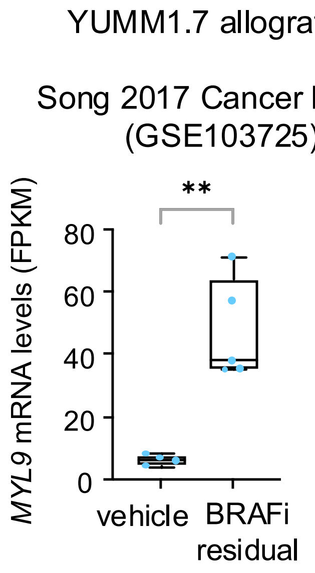
YUMM1.7 allografts
Song 2017 Cancer Discov
(GSE103725)
**
MYL9 mRNA levels (FPKM)
80
60
40
20
0
vehicle
BRAFi
residual

**Figure S3. NMII mRNA levels increase early on during adaptation to MAPK inhibition in vivo.** (**a**) Expression levels of NMII genes in published RNA-seq data of melanoma patient biopsies before and on BRAFi treatment (7,9). Heatmap shows $\log_2$ fold change of treated vs baseline biopsies. (**b**) *MYL9* expression levels in YUMM1.7 allografts treated or not with BRAFi from published RNA-seq data (7). FPKM, fragments per kilobase of transcript per million mapped reads. Veh = vehicle-treated tumours, BRAFi residual = maximally regressed, residual BRAFi-treated tumours. Boxplot: median (centre line); interquartile range (box); min-max (whiskers) and dots for individual data points.

Figure S4

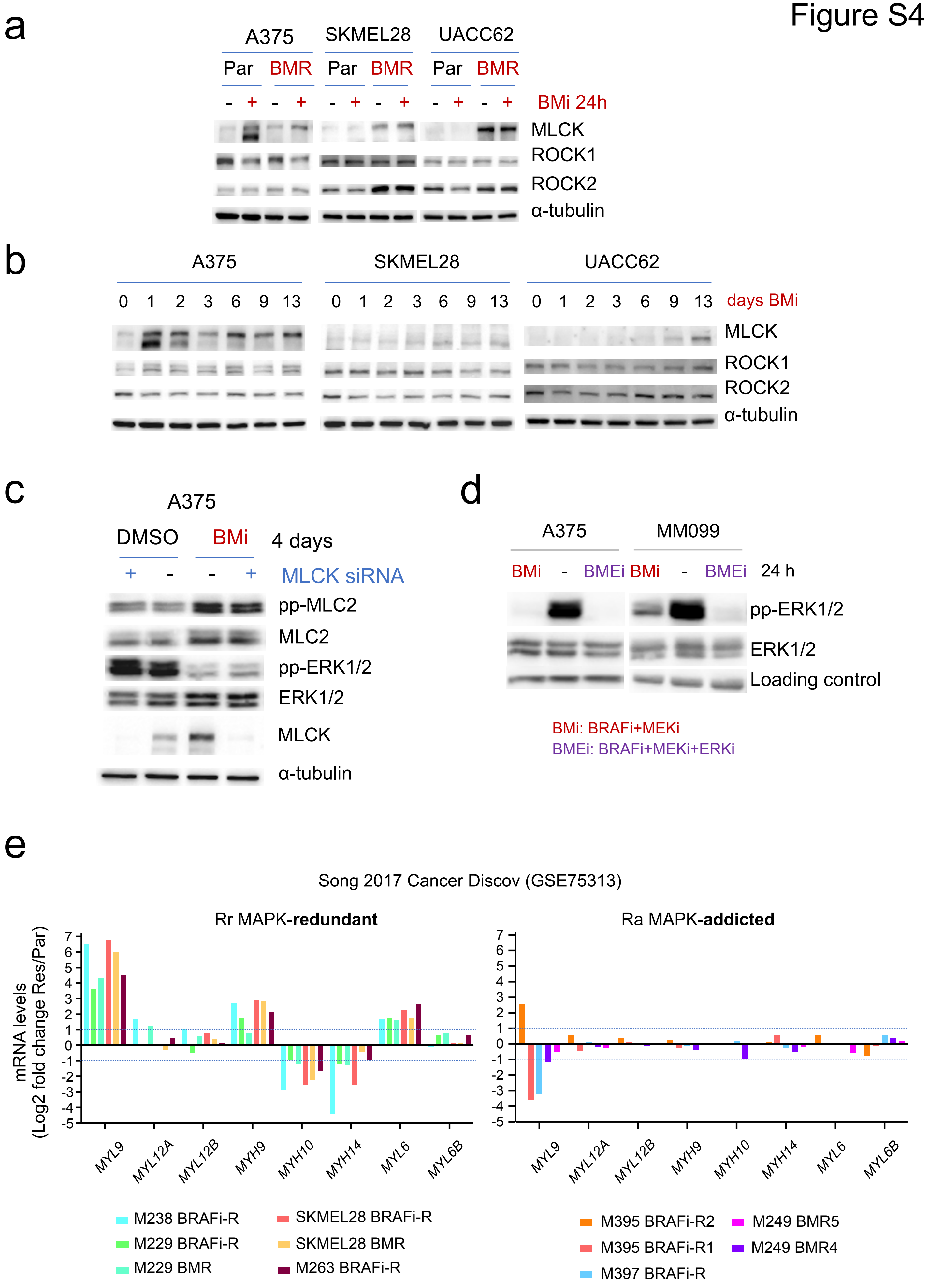
a
A375
SKMEL28
UACC62
Par BMR Par BMR Par BMR
- + - + - + - + - + - +
BMi 24h
MLCK
ROCK1
ROCK2
α-tubulin
b
A375
SKMEL28
UACC62
0 1 2 3 6 9 13
days BMi
MLCK
ROCK1
ROCK2
α-tubulin
c
A375
DMSO
BMi
4 days
+ - - +
MLCK siRNA
pp-MLC2
MLC2
pp-ERK1/2
ERK1/2
MLCK
α-tubulin
d
A375
MM099
BMi - BMEi BMi - BMEi 24 h
pp-ERK1/2
ERK1/2
Loading control
BMi: BRAFi+MEKi
BMEi: BRAFi+MEKi+ERKi
e
Song 2017 Cancer Discov (GSE75313)
Rr MAPK-**redundant**
Ra MAPK-**addicted**
mRNA levels (Log2 fold change Res/Par)
7 6 5 4 3 2 1 0 -1 -2 -3 -4 -5
MYL9 MYL12A MYL12B MYH9 MYH10 MYH14 MYL6 MYL6B
M238 BRAFi-R
M229 BRAFi-R
M229 BMR
SKMEL28 BRAFi-R
SKMEL28 BMR
M263 BRAFi-R
M395 BRAFi-R2
M395 BRAFi-R1
M397 BRAFi-R
M249 BMR5
M249 BMR4

**Figure S4. NMII overactivation during adaptation to BMi is controlled by ROCK rather than by ERK rebound.** (**a, b**) Immunoblots of indicated proteins from melanoma cell lines after treatments with BMi (125 nM BRAFi dabrafenib + 6.25 nM MEKi trametinib). Representative of 2 independent experiments. (**c**) Immunoblots of indicated proteins from A375 cells treated with BMi for 3 days and transfected with siRNA against MLCK. Non-targeting siRNA (-) was used as control. (**d**) Immunoblots of indicated proteins from melanoma cell lines after 24 h treatment with BMi (125 nM BRAFi dabrafenib + 6.25 nM MEKi trametinib) or BMEi (BMi plus 1 $\mu$M ERKi SCH772984). (**e**) Expression levels of NMII genes in published RNA-seq data of parental and BRAFi- or BMR melanoma cultures (7). mRNA levels are plotted as $\log_2$ fold change of resistant cells compared to parental cells. Dashed lines indicate changes larger or equal than two-fold. Resistant cells were classified as Rr (Resistant MAPK-redundant) and Ra (Resistant MAPK-addicted) as in the Song et al study (7). In the Song paper, BRAFi-resistant lines are called SDR (single drug resistant) and BMR lines DDR (double drug resistant).

Figure S5

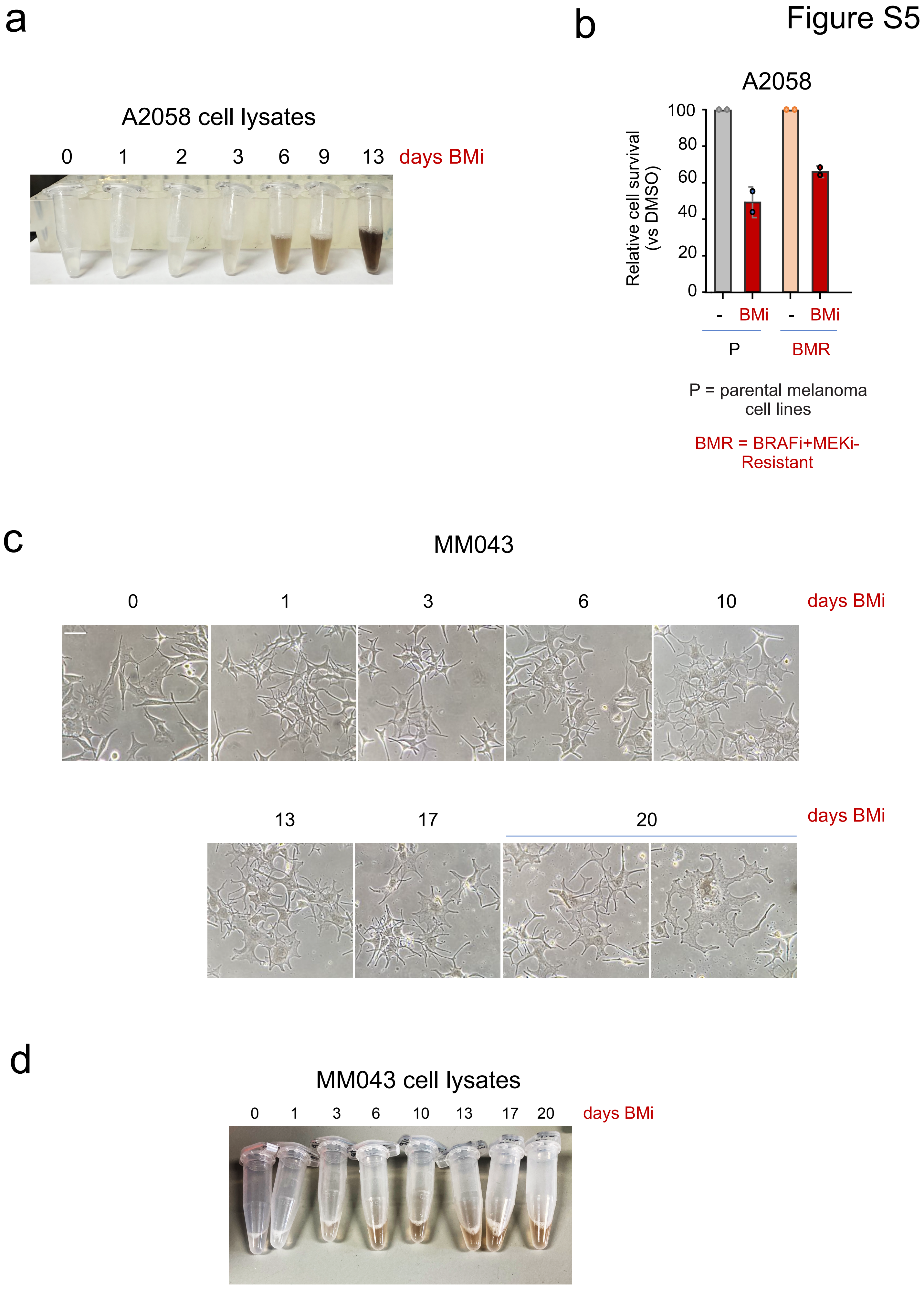
a
A2058 cell lysates
0 1 2 3 6 9 13 days BMi
b
A2058
Relative cell survival (vs DMSO)
100
80
60
40
20
0
- BMi - BMi
P
BMR
P = parental melanoma cell lines
BMR = BRAFi+MEKi-Resistant
c
MM043
0 1 3 6 10 days BMi
13 17 20 days BMi
d
MM043 cell lysates
0 1 3 6 10 13 17 20 days BMi

**Figure S5. NMII is not activated in melanomas with an hyperdifferentiated/pigmented phenotype during adaptation to BMi.** (**a**) Pictures of protein lysates from A2058 cells treated with BMi. (**b**) Relative cell survival by crystal violet staining of A2058 parental (P) and BMR cultures after BMi treatment for 3 days. n=2 independent experiments. (**c**) Representative phase-contrast images of MM043 treated with BMi. Note the presence of pigmented cells. Scale bar, 50 μm. (**d**) Pictures of protein lysates from MM043 cultures treated with BMi.

Figure S6

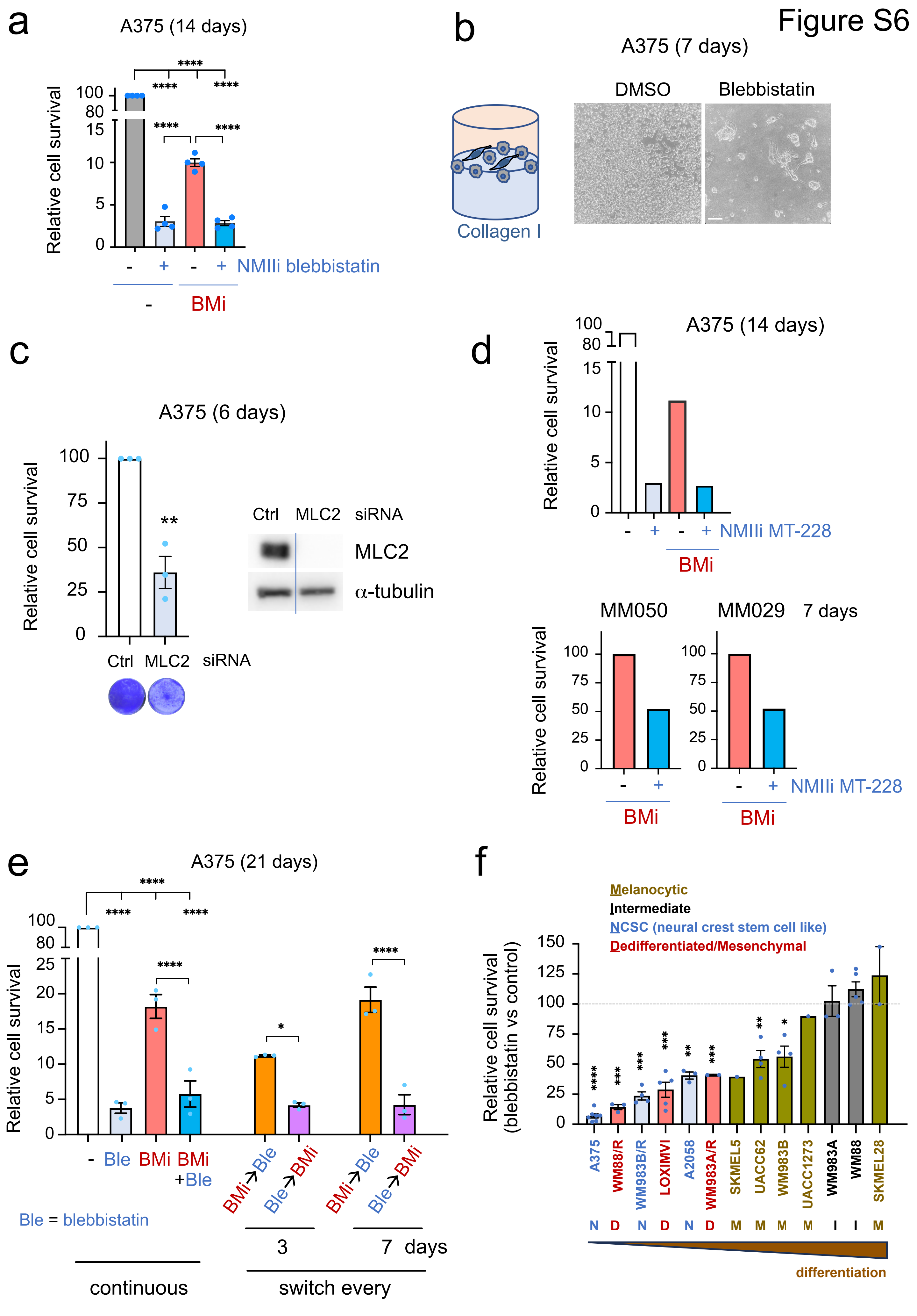
a
A375 (14 days)
Relative cell survival
100
80
15
10
5
0
****
****
****
****
****
- + - + NMIIi blebbistatin
-
BMi
b
A375 (7 days)
DMSO
Blebbistatin
Collagen I
c
A375 (6 days)
Relative cell survival
100
75
50
25
0
**
Ctrl MLC2 siRNA
Ctrl MLC2 siRNA
MLC2
α-tubulin
d
A375 (14 days)
Relative cell survival
100
80
15
10
5
0
- + - + NMIIi MT-228
BMi
MM050
MM029 7 days
Relative cell survival
100
75
50
25
0
- + - + NMIIi MT-228
BMi
BMi
e
A375 (21 days)
Relative cell survival
100
80
25
20
15
10
5
0
****
****
****
****
*
****
- Ble BMi BMi +Ble
BMi→Ble
Ble→BMi
BMi→Ble
Ble→BMi
Ble = blebbistatin
3
7 days
continuous
switch every
f
Melanocytic
Intermediate
NCSC (neural crest stem cell like)
Dedifferentiated/Mesenchymal
Relative cell survival (blebbistatin vs control)
150
125
100
75
50
25
0
****
***
***
***
**
***
**
*
A375
WM88/R
WM983B/R
LOXIMVI
A2058
WM983A/R
SKMEL5
UACC62
WM983B
UACC1273
WM983A
WM88
SKMEL28
N D N D N D M M M M I I M
differentiation

**Figure S6. NMII inhibition decreases tolerance of some melanomas to BRAFi+MEKi.** (**a**, **d**) Relative cell survival by crystal violet staining of melanoma cultures after treatment with BMi (125 nM BRAFi dabrafenib + 6.25 nM MEKi trametinib), 10 μM NMII inhibitor (blebbistatin or MT-228), or both (n = 3). (**b**) Representative phase-contrast images of A375 cells on top of a collagen I matrix and treated with 10 μM blebbistatin or vehicle DMSO for 7 days. Scale bar, 100 μm. (**c**) Relative cell survival by crystal violet staining of melanoma cultures after transfection with siRNA against MLC2 genes (*MYL9+MYL12A+MYL12B*) for 6 days. Representative crystal violet pictures and MLC2 immunoblots are shown. Line denotes non-contiguous lanes in the same gel/blot. (**e**) Relative cell survival by crystal violet staining of melanoma cultures after treatment with BMi (125 nM BRAFi dabrafenib + 6.25 nM MEKi trametinib), 10 μM blebbistatin (Ble) or both (n = 3). Cells were treated continuously for 21 days, or alternating treatments every 3 or 7 days. (**f**) Relative cell survival by crystal violet staining of melanoma cell lines treated with 10 μM blebbistatin for 7 days. Main cell phenotype is indicated. P-values by one-way ANOVA with Tukey post-hoc test (a, e), one sample t-test (f), t-test (c).
